# ILF and vReD pathways cooperate to control lysosomal transporter protein lifetimes

**DOI:** 10.1101/204396

**Authors:** Erin Kate McNally, Christopher Leonard Brett

## Abstract

Lysosomal nutrient transporter proteins move lumenal products of biomaterial catabolism to the cytoplasm for reuse by the cell. Two mechanisms control their lifetimes: the ILF (IntraLumenal Fragment) and vReD (Vacuole REcycling and Degradation) pathways. But it is not clear if they function independently. Using *S. cerevisiae* as a model, here we show that the ILF pathway mediates constitutive turnover of the lysine transporter Ypq1 and zinc transporter Cot1—known vReD client proteins—in vivo and in vitro. In contrast, the vReD pathway mediates constitutive degradation of the amino acid transporter Vba4. Activation of TOR with cycloheximide enhances their degradation by these pathways. However, misfolding by heat stress shunts all three into the ILF pathway. Thus, both pathways control individual transporter lifetimes, although only the ILF pathway mediates protein quality control. The pathway chosen depends on protein fate: degradation is imminent by the ILF pathway, whereas the vReD pathway permits reuse.

**HIGHLIGHTS and eTOC BLURB:**

- vReD and ILF pathways control Ypq1, Cot1 and Vba4 lifetimes
- ILF pathway constitutively degrades Ypq1 and Cot1, vReD degrades Vba4
- TOR activation stimulates protein degradation by both pathways
- ILF pathway clears all misfolded proteins for quality control

Two mechanisms degrade lysosome transporter proteins but it is not clear how each contributes to their lifetimes for organelle homeostasis or remodeling. Here McNally and Brett show that the transporters Cot1, Ypq1 and Vba4 can be selectively degraded by the ILF (IntraLumenal Fragment) pathway or vReD (Vacuole REcycling and Degradation) pathway depending on stimulus. However, only the ILF pathway mediates protein quality control.

## INTRODUCTION

Lysosomes are tasked with recycling biomaterials within all eukaryotic cells (*Perera and Zoncu, 2016*). In addition to being an important source of cellular nutrients, their activity is required to clear toxic aggregates or damaged proteins and organelles to prevent cell death. To function, lysosomes must perform three fundamental functions (*de Duve and Wattiaux, 1966; Luzio et al., 2007*): first they must fuse with membrane-encapsulated compartments containing biomaterials, which exposes them to lumenal acid hydrolases for catabolism–the second essential function of lysosomes. Finally, the products of degradation (lipids, sugars, amino acids, and other nutrients) are translocated across the lysosomal membrane by nutrient transporter proteins to the cytoplasm where they are re-used by the cell. In addition to acting as an important source of nutrients, lysosomes also store metals and other ions, and controlled release (or uptake) by lysosomal transporter proteins is critical for metabolism, signaling and programmed cell death (*Kroemer and Jäättelä, 2005*; *Xu and Ren, 2015*; *Lim and Zoncu, 2016*). Thus, lysosomal transporter protein expression and activity is critical for lysosome function, yet we know little about their lifetimes or how they are regulated.

Recently two mechanisms have been discovered that control polytopic protein lifetimes on the vacuolar lysosome (or vacuole) in the model organism *Saccharomyces cerevisiae*: The first, termed the vReD (vacuole membrane REcycling and Degradation) pathway, is essentially a mechanism to feed vacuolar polytopic proteins to the canonical MultiVesicular Body (MVB) pathway, which is responsible for surface transporter and receptor protein down-regulation (*Davies et al., 2009*; Figure 1A): In response to changes in substrate levels, the vacuolar amino acid transporter Ypq1 (*Li et al., 2015a*) or zinc transporter Cot1 (*Li et al., 2015b*) are ubiquitylated and sorted into areas of the vacuole membrane that form vesicles, which then supposedly fuse with endosomal compartments. Here they are thought to encounter ESCRTs (Endosomal Sorting Complexes Required for Transport) that sort and package them into intralumenal vesicle (ILVs) forming a MVB, which when mature, fuses with the vacuole to expose protein-laden ILVs to lumenal acid hydrolases for catabolism. Although the evidence presented confirms the existence of the vReD pathway, widespread use by vacuolar polytopic proteins has not been confirmed and its contributions to vacuole membrane homeostasis or remodeling remain elusive.

**Figure 1.**
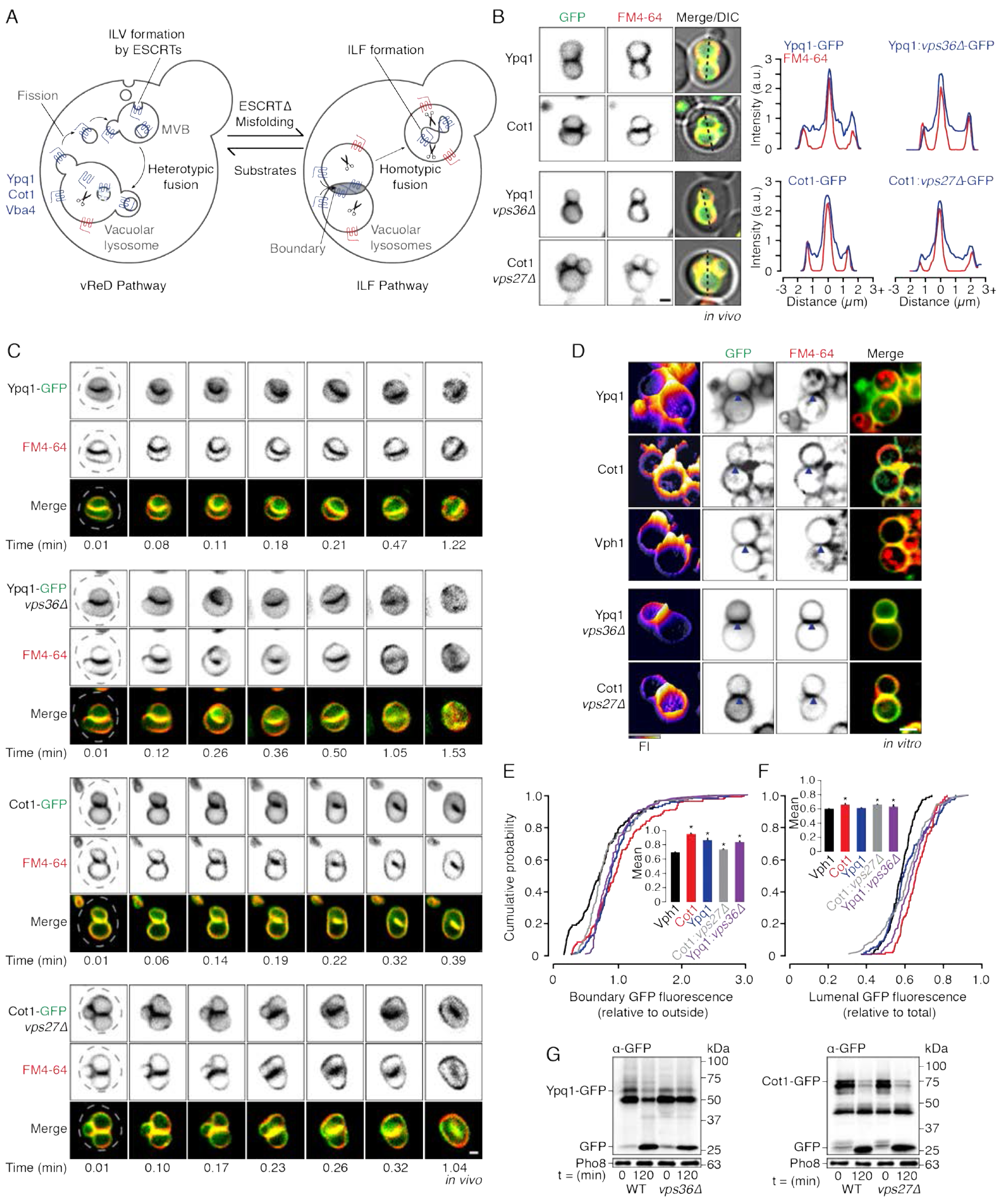
vReD client proteins Ypq1 and Cot1 are constitutively degraded by the ILF pathway. **(A)** Working model describing how lysosomal membrane proteins are selectively sorted for degradation by either the ILF or vReD pathways. **(B)** Fluorescence micrographs and line-scan analysis (dotted line) of live wild type or ESCRT deficient cells expressing Ypq1-GFP or Cot1-GFP stained with FM4-64 to label vacuole membranes. **(C)** Images from time-lapse videos of live yeast cells undergoing vacuole fusion expressing Ypq1-GFP or Cot1-GFP in wild type or ESCRT deficient cells stained with FM4-64. Dotted lines outline each cell as observed by DIC. **(D)** Micrographs of FM4-64-stained vacuoles isolated from wild type or ESCRT deficient cells expressing Ypq1-GFP, Cot1-GFP or Vph1-GFP observed under standard fusion conditions. Closed arrowheads indicate boundary membranes containing GFP fluorescence. GFP fluorescence intensity profiles are shown (left panels). Using micrographic data presented in D, we generated cumulative probability plots for GFP fluorescence intensity within the boundary membrane **(E)** or vacuole lumen (F; n > 102). Means (± S.E.M.) are plotted and compared to values obtained for Vph1-GFP (inserts). *P < 0.05. Western blot analysis of Ypq1-GFP **(G)** or Cot1-GFP (H) degradation before (0) or after (120 min) fusion of vacuoles isolated from wild type (WT) or ESCRT deficient cells. Alkaline phosphatase (Pho8) is shown as a load control. Scale bars, 1 μm (in vivo) or 2 μm (in vitro).

The second, termed the IntraLumenal Fragment (ILF) pathway, is an ESCRT-independent mechanism that relies on vacuole membrane fusion machinery for protein sorting and internalization (*McNally et al., 2017*; Figure 1A): In response to substrates, protein misfolding or TOR-signaling, vacuolar polytopic proteins such as the V-ATPase (Vph1), metal transporters (Fth1, Fet5), ABC (ATP-Binding Cassette) transporters (Ybt1, Ycf1) and the lipid transporter Ncr1 are selectively sorted into an area of membrane, called the boundary, encircled by the fusion machinery that assembles into a ring at contact sites between apposing organelles prior to homotypic fusion. Upon lipid bilayer scission at the ring, the boundary membranes merge and are internalized within the lumen of single organelle as a protein-laden ILF that is catabolized by hydrolases (*Wang et al., 2002*; *Mattie et al., 2017*). Although the precise molecular mechanism responsible for protein sorting is unknown, this process requires activation of the Rab GTPase Ypt7, its downstream effector tethering complex HOPS (HOmotypic fusion and Protein Sorting) and formation of the ILF requires SNAREs (Soluble NSF Attachment protein REceptors; *Mattie et al., 2017; McNally et al., 2017*). While this pathway is crucial for remodeling the vacuolar membrane proteome, how it contributes to vacuole physiology or cellular metabolism remains understudied.

Although these pathways seem mutually exclusive, the outcome is equal: To topologically accommodate degradation by lumenal hydrolases, vacuolar polytopic proteins are either sorted and packaged into ILVs by ESCRTs at endosomes by the vReD pathway, or into ILFs – which are essentially large ILVs – by the fusion machinery on vacuoles by the ILF pathway (Figure 1A). But it is unclear why vacuolar polytopic proteins take different routes. Furthermore, Emr and colleagues reported that they were unable to visualize entry of some ILF client proteins into the vReD pathway (e.g. Vph1, Fth1; *Li et al., 2015a*), suggesting that protein degradation by these pathways is mutually exclusive and possibly dependent on protein identity. However, we found that the vReD client protein Cot1, a vacuolar zinc transporter, is also degraded by the ILF pathway (*Li et al., 2015b*; *McNally et al. 2017*). This preliminary work raises many important questions that must be answered to comprehensively understand vacuole transporter regulation and its contribution to cell biology. For example, can both pathways degrade the same client proteins? How is a protein selectively sorted into one pathway or the other? Does each pathway contribute to vacuole membrane homeostasis or protein quality control? Does each mediate vacuolar lysosome membrane remodeling in response to signaling responsible for cellular metabolism or aging?

## RESULTS

### vReD client proteins Cot1 and Ypq1 are constitutively degraded by the ILF pathway

To better resolve how vReD and ILF pathways contribute to vacuole membrane protein turnover, we first determined if the vReD client protein Ypq1 (*Li et al., 2015a*) is also degraded by the ILF pathway. Recently, Emr and colleagues suggested that Ypq1 is exclusively degraded by the vReD pathway (*Zhu et al., 2017*). This conclusion is based on primarily studying yeast cells that contain a single vacuole, eliminating the possibility of detecting the ILF pathway, as it requires fusion of two organelles. However, most cells within an actively growing culture contain two or more vacuoles (*Li and Kane, 2009*). Thus, to resolve this issue, we simply imaged live yeast cultures under standard growth conditions and examined cells with multiple vacuoles. Similar to Cot1, we find that GFP-tagged Ypq1 is present in the boundary membranes of docked organelles (Figure 1B). Using HILO microscopy, we then recorded homotypic vacuole fusion events in live cells and found that Ypq1-GFP embedded in the boundary membrane is internalized within the lumen upon the completion of lipid bilayer merger (Figure 1C; Video 1). Closer examination of micrographs presented in the three publications describing the vReD pathway reveal that fluorescent protein-tagged Ypq1 and Cot1 also accumulate within the boundary membrane between docked vacuoles when images of untreated yeast cells containing two or more vacuoles are shown (*Li et al., 2015a Figures 2A, 2C and 4I; Li et al., 2015b Figures 5D, 6C and 7C; Zhu et al., 2017 Figure 5A*). Thus, our results are consistent with these previous reports, and suggest that, like Cot1-GFP, Ypq1-GFP is constitutively sorted for degradation by the ILF pathway.

When degraded by the vReD pathway, Ypq1-GFP is recognized, sorted and packaged for degradation by the ESCRT machinery (*Li et al., 2015a*). Thus, it is possible that ESCRTs may sort Ypq1-GFP into the boundary as well. To test this hypothesis, we deleted VPS36, a gene encoding a subunit of ESCRT-I (*Henne et al., 2011*) to disable the ESCRT machinery and found that it had no effect on Ypq1-GFP sorting into the ILF pathway (Figure 1B and C; Video 2). We made similar observations for Cot1-GFP when VPS27, a gene encoding a component of ESCRT-0 (*Katzmann et al., 2003*) was deleted (Figure 1B and C; Videos 3 and 4). This finding is consistent with our previous work confirming that protein degradation by the ILF pathway is ESCRT-independent (*McNally et al., 2017*) and eliminates the possibility that the vReD pathway is responsible for the observed sorting phenotypes.

The machinery necessary for protein degradation by the ILF pathway co-purifies with isolated organelles permitting study in vitro (*McNally et al., 2017*). Thus, we next confirmed that Ypq1-GFP sorting into the ILF pathway observed in vivo also persists in vitro (Figure 1D). We measured the GFP fluorescence intensity at these boundary membranes and found that, like Cot1-GFP, Ypq1-GFP was enriched at these sites (Figure 1E), as compared to GFP-tagged Vph1, the stalk domain of the vacuolar V-type H+-ATPase, which is uniformly distributed on vacuole membranes during fusion (*Wang et al., 2002; McNally et al., 2017*). Furthermore, protein sorting was unaffected by deleting components of the ESCRT machinery, confirming that entry of Ypq1 and Cot1 into the ILF pathway is independent of ESCRTs and the vReD pathway. To confirm that Ypq1-GFP was internalized during the fusion reaction in vitro, we sought to determine if Ypq1-GFP appeared within the lumen over time (Figure 1F). As expected, we found that Ypq1-GFP and Cot1-GFP accumulated within the lumen, similar to Vph1-GFP under fusogenic conditions. Here, these proteins should be exposed to acid hydrolases for degradation (*Luzio et al., 2007*). Thus, we next conducted western blot analysis to assess protein degradation and found that more GFP was cleaved from Ypq1 and Cot1 after lysosome fusion was stimulated in vitro (Figure 1G). Importantly, protein degradation continued to occur in the absence of ESCRT activity. In all, these results reveal that both known vReD client proteins, Ypq1 and Cot1, are constitutively turned over by the ILF pathway.

### Vba4, a vacuolar amino acid transporter, is constitutively degraded by the vReD pathway

At this point, it seems that constitutive turnover of vacuolar polytopic proteins exclusively relies on the ILF pathway (see *McNally et al., 2017*). However, when conducting studies to identify additional ILF client proteins, we discovered that GFP-tagged Vba4, a vacuole amino acid transporter (*Kawano-Kawada et al., 2015*), was excluded from boundary membranes between docked vacuoles in living cells (Figures 2A and B; Video 5). We quantified this phenotype using still images (Figure 2C) and confirmed that it is prevalent within the population of cells analyzed (Figure 2D). This sorting phenotype (boundary exclusion) is similar to other resident transporter proteins that were reported to avoid the ILF pathway and are spared during homotypic vacuole fusion (e.g. GFP-tagged Fet5, Ypt1, Ycf1 or Ncr1; *McNally et al., 2017*). However, unlike these other transporter proteins, Vba4-GFP also appeared on puncta adjacent to the vacuole membranes (Figures 2A) within most cells analyzed (Figure 2D). Importantly, these puncta resembled Cot1-GFP or Ypq1-GFP -positive structures that form to mediate their degradation by the vReD pathway in response to changes in substrate levels (Figures 2A, C and D; *Li et al., 2015a; Li et al., 2015b*). Consistent with this interpretation, in the presence of ZnCl_2_ Cot1-GFP was also excluded from boundary membranes and appeared on puncta adjacent to the vacuole membrane (Figure 2A), suggesting that it is redirected from the ILF pathway to the vReD pathway. However, despite the appearance of Ypq1-GFP on puncta, the GFP-tagged protein continued to be present in boundary membranes after lysine withdrawal (Figures 2A, C and D), suggesting that Ypq1-GFP substrate-induced degradation is not exclusively dependent on the vReD pathway. In any case, together these results show that Vba4 avoids the ILF pathway and rather is a genuine vReD client protein, the first shown to be constitutively degraded by this pathway.

**Figure 2.**
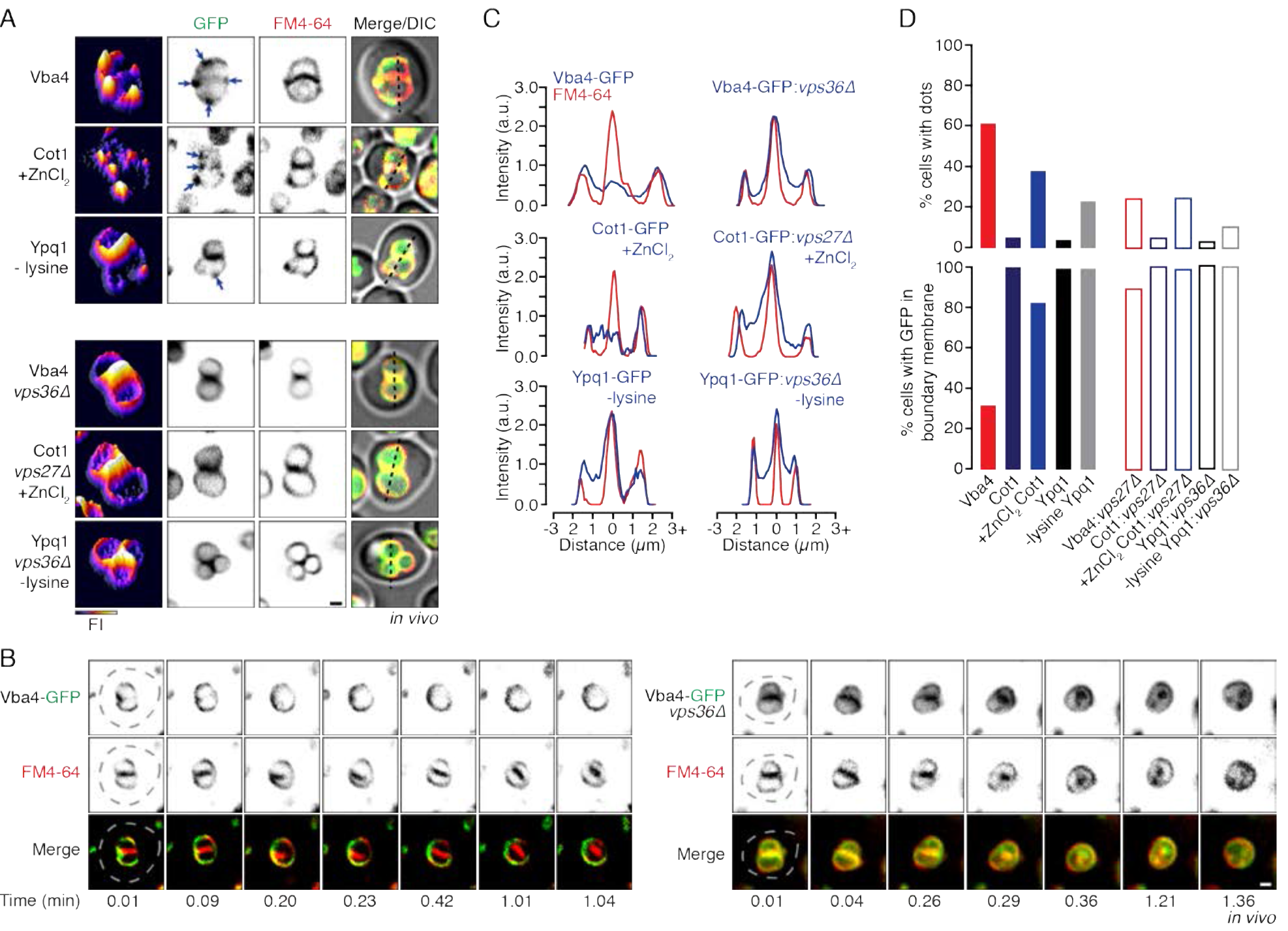
Vba4 is constitutively degraded by the vReD pathway in live cells. **(A)** Fluorescence micrographs of live wild type or ESCRT deficient yeast cells expressing Vba4-GFP, Cot1-GFP, or Ypq1-GFP stained with FM4-64 to label vacuole membranes. Addition of 2 mM ZnCl_2_ or lysine withdrawal was used to drive Cot1-GFP or Ypq1-GFP into the vReD pathway, respectively. Arrows highlight puncta (dots) adjacent to vacuole membranes. GFP fluorescence intensity profiles are shown (left panels). **(B)** Images from time-lapse videos of homotypic vacuole fusion events within live wild type or *vps36Δ* cells expressing Vba4-GFP stained with FM4-64. Dotted lines outline each cell as observed by DIC. **(C)** Fluorescence line-scan analysis of images shown in A (dotted lines). **(D)** Using micrographic data shown in A and Figure 1B, we calculated percentage of cells with GFP fluorescence present within the boundary membrane (bottom) and the percentage of cells with dots adjacent to vacuole membranes (top; n ≥ 155). Scale bars, 1 μm.

To further study protein degradation by these two pathways, we purified vacuoles from these strains and found that puncta form adjacent to isolated vacuoles when we monitored Vba4-GFP or Cot1-GFP (with ZnCl_2_) under fusogenic conditions over time (Figure 3A). This suggests that the vReD machinery responsible for protein sorting and packaging also co-purifies with organelles permitting further study in vitro. Notably, Ypq1-GFP did not appear on puncta in vitro (Figure 1D) despite the absence of lysine from the reaction buffer, suggesting that the machinery responsible for sensing extracellular lysine withdrawal to signal Ypq1-GFP degradation by the vReD pathway does not co-purify with vacuoles. Thus, we focused on studying Vba4-GFP and Cot1-GFP and found that both proteins continued to be excluded from boundaries between docked organelles (Figure 3C) and appeared on puncta (Figure 3B) over time, confirming that their sorting phenotypes were recapitulated in vitro. Importantly, concentrations of ZnCl_2_ used to stimulate Cot1-GFP sorting into the vReD pathway do not block the in vitro fusion reaction (Figure S1), suggesting the membrane fusion machinery and ILF pathway is intact but does not contribute to Cot1-GFP degradation under these conditions.

**Figure 3.**
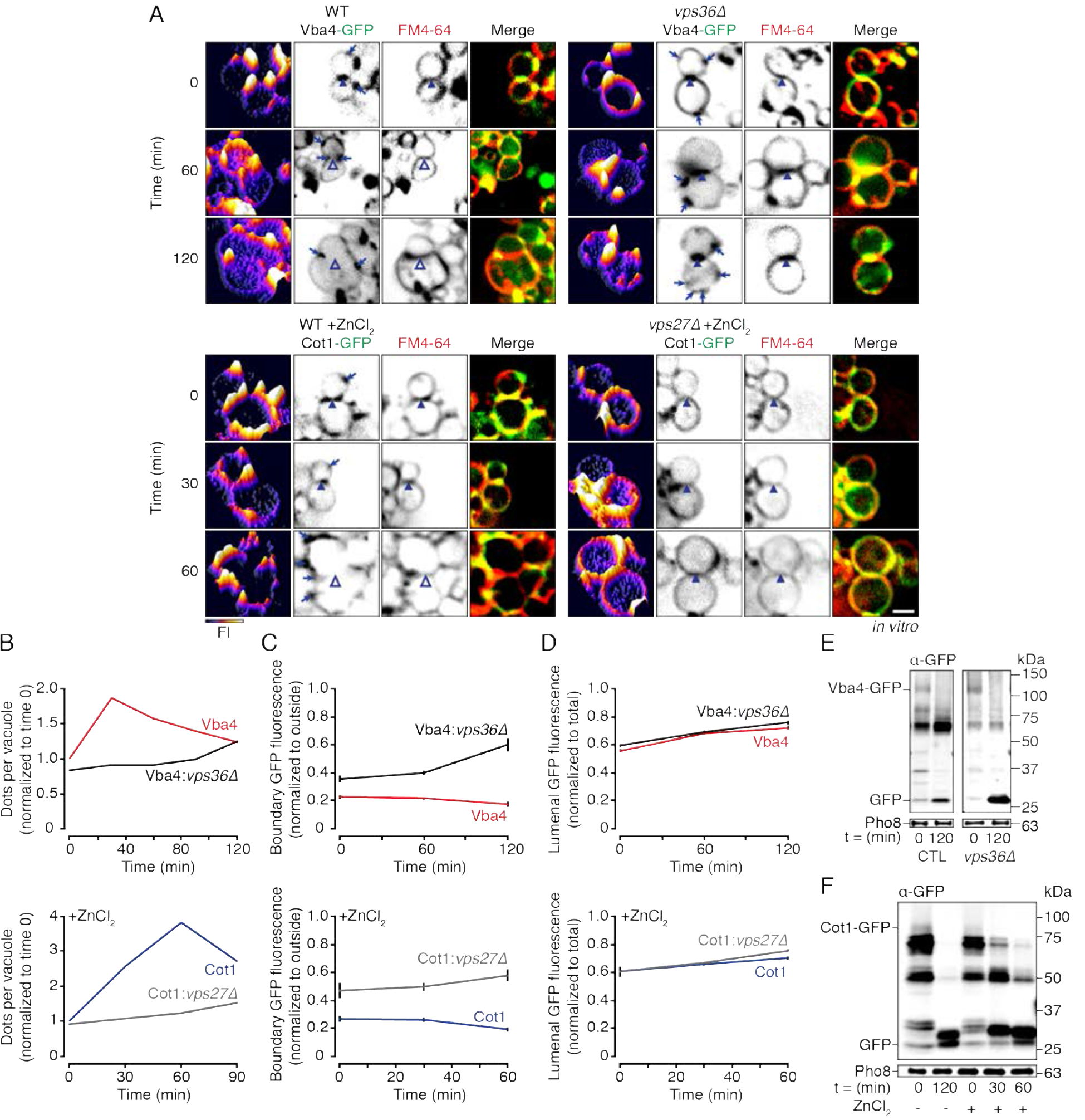
Vba4-GFP and Cot1-GFP are degraded by the vReD pathway in vitro. Fluorescence micrographs of vacuoles isolated from wild type or ESCRT deficient cells expressing Vba4-GFP or Cot1-GFP stained with FM4-64 observed under standard fusion conditions over time. Vacuoles were pre-treated with 25 μM ZnCl_2_ for 5 minutes to drive Cot1-GFP into the vReD pathway. Arrows indicate puncta adjacent to vacuole membranes. Boundary membranes containing (closed arrowheads) or lacking (open arrowheads) GFP fluorescence are indicated. GFP fluorescence intensity profiles are shown (left panel). Scale bar, 2 μm. Using micrographic data shown in **A**, we calculated the mean (± S.E.M.) number of dots per vacuole (normalized to wild type at 0 min; B) or GFP fluorescence intensity in the boundary membrane **(C)** or vacuole lumen **(D)** over time (n ≥ 123). Western blot analysis of Vba4-GFP **(E)** or Cot1-GFP **(F)** degradation before (0) or after (30, 60, 120 min) fusion of vacuoles isolated from wild type or *vps36*Δ cells. Cot1-GFP degradation was also assessed with or without 25 μM ZnCl_2_. Pho8 is shown as a load control.

When counting GFP-positive puncta that formed over time in vitro, we noticed that the number of puncta initially increased, peaked at 30 minutes for Vba4-GFP or 60 minutes for Cot1-GFP and then decreased afterwards, suggesting that these compartments were generated and then consumed during the reaction (Figure 3B). This result is consistent with Cot1-GFP and Vba4-GFP being first sorted by ESCRTs (production of puncta) which are later delivered to the vacuole lumen for degradation (consumption of puncta; *Li et al., 2015a*). To test this hypothesis, we measured the fluorescence intensity of Vba4-GFP and Cot1-GFP (with ZnCl_2_) within the lumen over time (Figure 3D). As expected, both GFP-tagged proteins accumulated within the lumen, signifying that they were exposed to acid hydrolases for degradation. To confirm, we assessed GFP-cleavage from Vba4 or Cot1 (with ZnCl_2_) by western blot (Figures 3E and F) and found that both proteins were indeed degraded over time in vitro. As Vba4-GFP is excluded from boundary membranes, but rather appears on vesicles that are consumed over time under standard fusion conditions, we conclude that Vba4-GFP is constitutively delivered to the vacuole for degradation by the vReD pathway, whereas in response to changing substrate levels, Cot1-GFP is rerouted from the ILF to the vReD pathway for degradation.

### The ILF pathway compensates for loss of vReD function

The vReD pathway requires ESCRTs to recognize, sort and package proteins for degradation. To confirm that the vReD pathway is responsible for constitutive Vba4-GFP degradation, we deleted VPS36 to block ESCRT function and expected Vba4-GFP to accumulate in puncta (as this stage of the vReD pathway is supposedly ESCRT-independent) but it should not be delivered to the vacuole lumen, as ILV formation is ESCRT-dependent (*Li et al., 2015a*). However, we found that knocking out VPS36 caused Vba4-GFP to accumulate within the boundary membrane at the interface between docked vacuoles (Figures 2A–C) in nearly all cells examined (Figure 2D), and Vba4-GFP was internalized within an ILF upon homotypic vacuole fusion within living cells (Figure 2B; Video 6). We made similar observations in vitro (Figures 3A) whereby the intensity of Vba4-GFP fluorescence within the boundary membrane increased over time (Figure 3C), suggesting that it was now sorted into the ILF pathway.

Moreover, Vba4-GFP-positive puncta did not efficiently form in vivo (Figure 2D) or in vitro (Figure 3B), although Vba4-GFP continued to accumulate in the lumen over time (Figure 3D). Consistent with this finding, Vba4-GFP continued to be degraded in the absence of VPS36 (Figure 3E). We made similar observations in vitro for Cot1-GFP, whereby ZnCl_2_ normally triggers Cot1-GFP degradation by the vReD pathway, but when ESCRT function is disrupted, it resorts to the ILF pathway (Figures 2 and 3). All things considered, we conclude that vReD-mediated protein degradation is rerouted to the ILF pathway when ESCRT function is impaired.

### Protein degradation by the vReD pathway can be stimulated by TOR

TOR (Target Of Rapamycin) is a serine/threonine kinase important for regulating cellular metabolism during growth, stress or aging (*Dunlop and Tee, 2009; Laplante and Sabatini, 2012; Perera and Zoncu, 2016*). To presumably regulate availability of cytoplasmic amino acid levels, the ILF and MVB pathways degrade vacuolar and surface amino acid transporters, respectively, in response to TOR signaling upon the addition of cycloheximide (CHX; *MacGurn et al., 2011; McNally et al., 2017*). However, it is not known if vReD also mediates vacuolar transporter protein degradation in response to TOR signaling. To test this hypothesis, we treated cells with CHX and examined the distribution of Vba4-GFP, the only client protein that is constitutively degraded by the vReD pathway. We find that Vba4-GFP continues to be excluded from boundary membranes (Figures 4A and B; Video 7) within the cell population (Figure 4C) but more puncta appear adjacent to vacuoles in the presence of CHX (Figure 4C). In addition, we find that CHX increases cleavage of GFP from Vba4 (Figure 4D), indicating that degradation of Vba4-GFP by the vReD pathway is enhanced. Importantly, treatment with rapamycin, a TOR signaling inhibitor, blocked this effect (Figures 4A and C), confirming that TOR activation was required. We made similar observations in vitro (Figures 4E–G) and found that more Vba4-GFP accumulated in the vacuole lumen in the presence of CHX (Figure 4H). Consistent with this observation, GFP cleavage from Vba4 was enhanced in the presence of CHX (Figure 4I). Importantly, the observed increase in Vba4-GFP degradation was blocked by pretreating isolated organelles with rapamycin (Figure 4I). Thus, we conclude that TOR activation enhances Vba4-GFP degradation by the vReD pathway.

**Figure 4.**
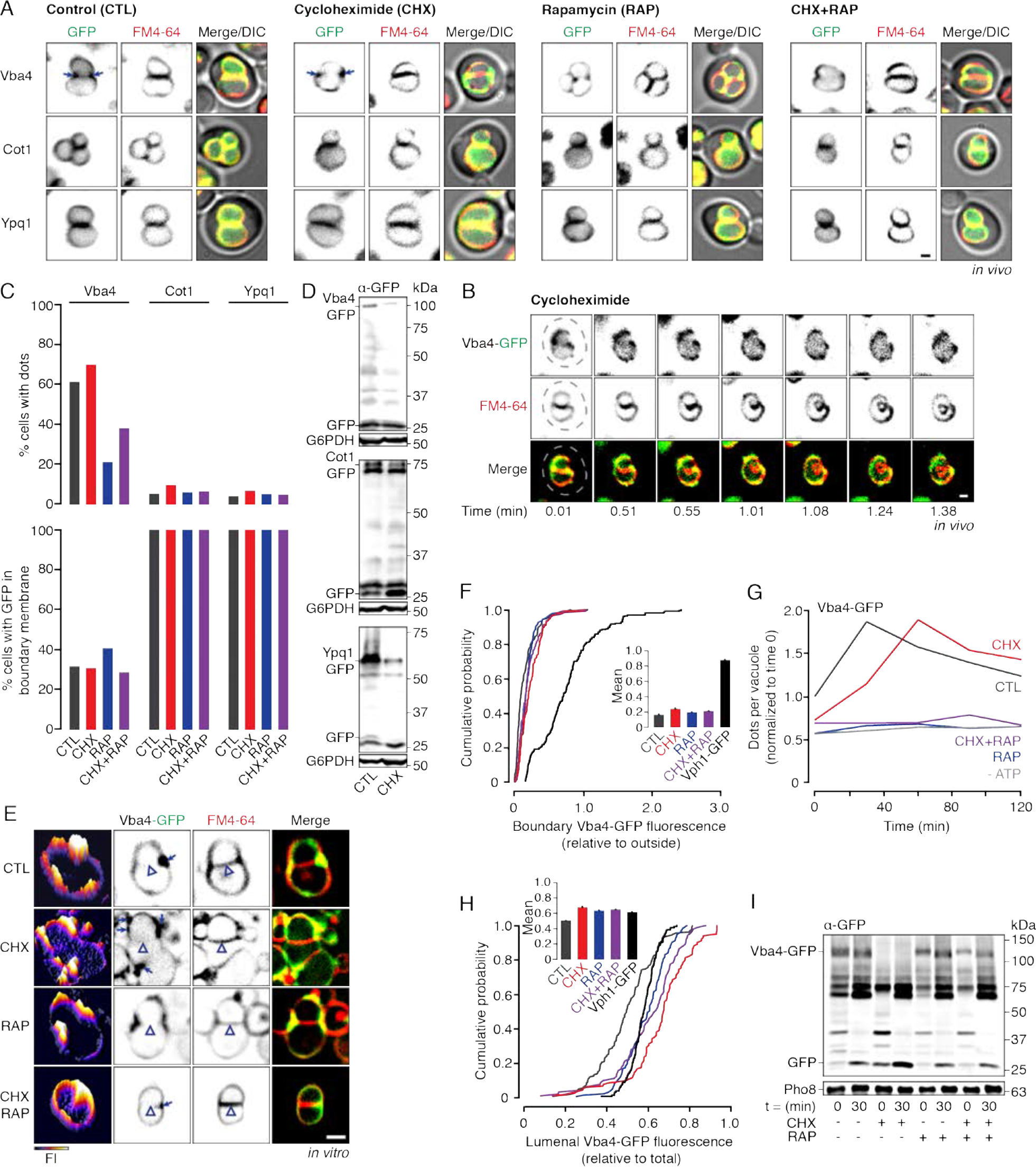
Protein degradation by the vReD pathway is stimulated in response to TOR signaling. **(A)** Fluorescence micrographs of live wild type cells expressing Vba4-GFP, Cot1-GFP or Ypq1-GFP stained with FM4-64 to label vacuole membranes under control conditions (CTL) or after incubation with cycloheximide (CHX), rapamycin (RAP) or both. Arrows highlight puncta adjacent to vacuole membranes. **(B)** Images from a time-lapse video of a vacuole fusion event within a live wild type cell expressing Vba4-GFP after CHX treatment. FM4-64 was used to label vacuole membranes. Dotted line outlines the yeast cell as observed by DIC. **(C)** Using micrographic data shown in **A**, we calculated percentage of cells with GFP fluorescence present within the boundary membrane (bottom) and the percentage of cells with dots adjacent to vacuole membranes (top; n ≥ 204). (D) Western blot analysis of Vba4-GFP, Cot1-GFP or Ypq1-GFP degradation within wild type cells in the presence or absence of CHX. Glucose-6-phosphate dehydrogenase (G6PDH) is shown as a load control. **(E)** Fluorescence micrographs of vacuoles isolated from wild type cells expressing Vba4-GFP stained with FM4-64 observed under standard fusion conditions (CTL) or after treatment with CHX, RAP or both. Boundary membranes lacking GFP fluorescence are indicated by open arrowheads. Arrows highlight puncta adjacent to vacuole membranes. GFP fluorescence profiles are shown (left panels). Using micrographic data presented in E, we generated cumulative probability curves of GFP fluorescence intensity within the boundary membrane **(F)** or vacuole lumen **(H)**. Means ± S.E.M. are shown (inserts; n ≥ 359 vacuoles). **(G)** Based on micrographic data shown in E, we calculated the mean (α S.E.M.) number of dots per vacuole (normalized to CTL at 0 min) over time (n ≥ 359 vacuoles). **(I)** Western blot analysis of Vba4-GFP degradation before (0) or after (30 min) fusion of vacuoles isolated from wild type cells in the presence or absence of CHX, RAP or both. Pho8 is shown as a load control. Scale bars, 1 μm (in vivo) and 2 μm (in vitro).

Can other proteins get shunted into the vReD pathway in response to TOR signaling? Although Ypq1-GFP and Cot1-GFP are constitutively degraded by the ILF pathway (see Figure 1), they are rerouted into the vReD pathway in response to changing substrate levels (Figure 2). Thus, it is possible that TOR activation by CHX may also drive Ypq1-GFP or Cot1-GFP into the vReD pathway for degradation. To answer this question, we treated cells expressing either protein with CHX and examined the effects on their cellular membrane distribution (Figure 4A–C). Both proteins continued to be present in boundary membranes in all cells, and neither appeared on puncta, suggesting that CHX did not reroute Ypq1-GFP or Cot1-GFP from the ILF pathway to the vReD pathway. Cleavage of GFP from both proteins was enhanced in the presence of CHX, suggesting TOR activation stimulates their degradation by the ILF pathway (Figure 4D). These results were confirmed in vitro (Figure S2A), whereby CHX enhanced sorting into the boundary membrane (Figure S2B), increased lumenal delivery (Figure S2C) and increased degradation (Figure S2D) of Ypq1-GFP and Cot1-GFP. Furthermore, protein degradation was blocked by fusion inhibitors (Figure S2E), confirming that enhanced degradation of Ypq1-GFP and Cot1-GFP by TOR activation was mediated by the ILF pathway instead of the vReD pathway.

### The ILF pathway exclusively mediates vacuolar polytopic protein quality control

To prevent cellular proteotoxicity and maintain proteostasis, misfolded or damaged proteins are selectively cleared by multiple degradation pathways, such as the proteasome or chaperone-mediate autophagy for cytoplasmic proteins (*Cuervo and Dice, 2000; Kaushik et al., 2011*), the ESCRT pathway for some surface polytopic proteins (*Babst, 2014*), and the ILF pathway for vacuolar polytopic proteins including Cot1 (*McNally et al., 2017*). Although its contribution to protein quality control has not been assessed, the vReD pathway relies on similar ESCRT machinery that clears misfolded surface proteins (*Raiborg and Stenmark, 2009; Li et al., 2015a*). Thus, it is possible that, like the ILF pathway, the vReD pathway may clear some misfolded vacuolar transporter proteins. To test this hypothesis, we subjected cells to heat stress to induce protein misfolding (*Keener and Babst, 2013; McNally et al., 2017*) and monitored changes in the membrane distributions Vba4-GFP, Ypq1-GFP and Cot1-GFP within living cells (Figures 5A and B). We find that Vba4-GFP is sorted into the boundary membrane between docked organelles in response to heat stress in nearly all cells analyzed (Figure 5C), similar to Ypq1-GFP, as well as Cot1-GFP (as previously reported; *McNally et al., 2017*). Because Vba4-GFP is normally excluded from boundary membranes, we also used HILO microscopy to confirm that it was indeed internalized within an ILF upon homotypic lysosome fusion after heat stress within living cells (Figure 5D; Video 8). We made similar observations when VPS36 or VPS27 were deleted, confirming that sorting of misfolded proteins into the boundary in response to heat stress was ESCRT-independent (Figures 5A–C). When we examined the effect of heat stress on protein degradation (Figure 5E), we find that cleavage of GFP from Ypq1 and Cot1 was enhanced, as expected. However, cleavage of GFP from Vba4 was unaffected, suggesting that Vba4-GFP degradation occurred at the same rate after heat shock, but this protein was rerouted from the vReD to ILF pathway for lumenal delivery.

**Figure 5.**
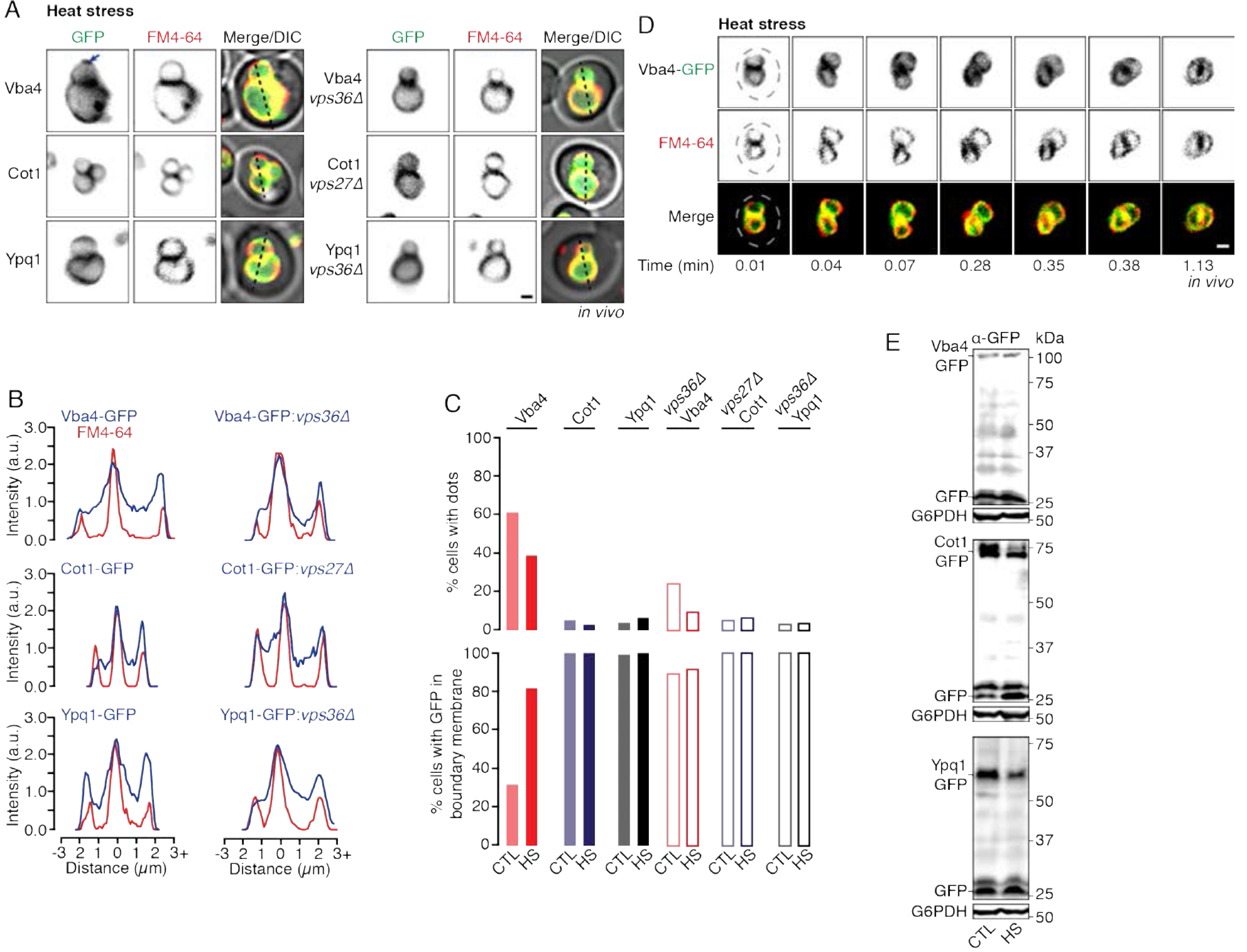
The ILF pathway degrades misfolded lysosomal polytopic proteins within live cells. **(A)** Fluorescence micrographs of live wild type or ESCRT deficient cells expressing Vba4-GFP, Cot1-GFP or Ypq1-GFP after heat stress. Cells were stained with FM4-64 to label vacuole membranes. Arrow highlights puncta adjacent to vacuole membranes. **(B)** Fluorescence line-scan analysis of images shown in **A** (dotted lines). **(C)** Using micrographic data shown in **A** and Figures 1A and 2A, we calculated percentage of cells with GFP fluorescence present within the boundary membrane (bottom) and the percentage of cells with dots adjacent to vacuole membranes under control conditions (CTL) or after heat stress (HS; n ≥148). (D) Images from a time-lapse video of a vacuole fusion event within a live wild type cell expressing Vba4-GFP after heat stress. FM4-64 was used to label vacuole membranes. Dotted line outlines the cell as observed by DIC. **(E)** Western blot analysis of Vba4-GFP, Cot1-GFP or Ypq1-GFP degradation within wild type cells before or after heat stress. Glucose-6-phosphate dehydrogenase (G6PDH) is shown as a load control. Scale bars, 1 μm.

We replicated these results in vitro (Figure 6A), whereby heat stress stimulated sorting of Vba4-GFP into the boundary membrane (Figure 6B), enhanced accumulation within vacuole lumen (Figure 6C) and GFP-cleavage by western blot (Figure 6D). Sorting, internalization and degradation of Cot1-GFP and Ypq1-GFP, however, were minimally affected by heat stress. These results confirm that protein misfolding by heat stress stimulates degradation of Vba4 by rerouting it from the vReD to the ILF pathway, whereas constitutive turnover of Cot1 and Ypq1 by the ILF pathway seems to be already elevated and thus unaffected by heat stress. These latter proteins are perhaps more susceptible to damage by lumenal hydrolases or are less stable and thus have shorter half-lives than Vba4 under standard conditions.

**Figure 6.**
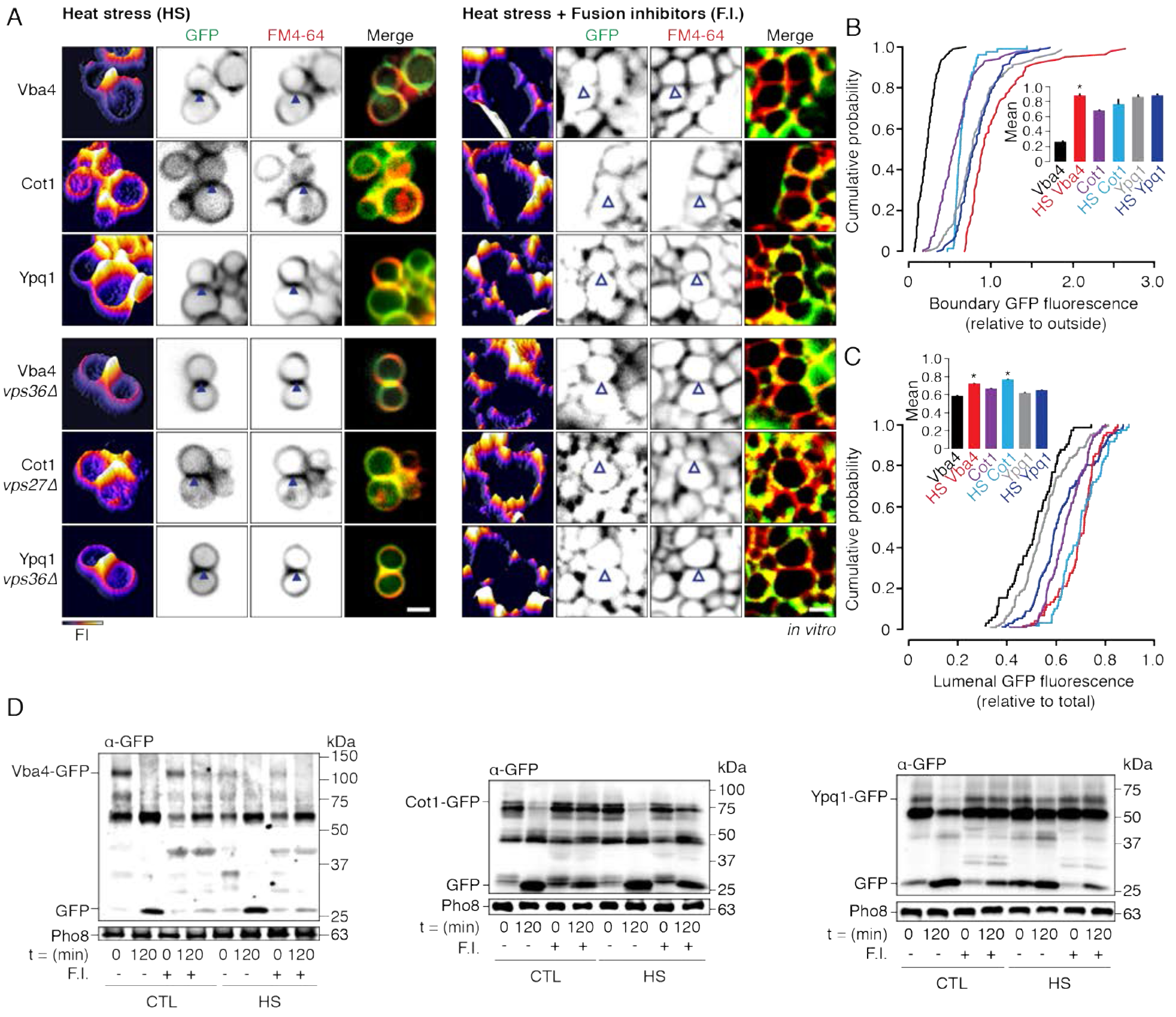
The ILF pathway degrades polytopic proteins when misfolded by heat stress in vitro. **(A)** Fluorescence micrographs of vacuoles isolated from wild type or ESCRT deficient cells expressing Vba4-GFP, Cot1-GFP or Ypq1-GFP after treatment with heat stress (HS) in the absence or presence of the fusion inhibitors rGdi1 and rGyp1-46 (F.I.). FM4-64 was used to label vacuole membranes. Boundary membranes containing (closed arrowhead) or lacking (open arrowhead) GFP fluorescence are indicated. GFP fluorescence profiles are shown (left panel). Scale bar, 2 μm. Using micrographic data presented in **E**, we generated cumulative probability curves of GFP fluorescence intensity within the boundary membrane **(B)** or vacuole lumen **(C)**. Means (± S.E.M.) are plotted and values obtained in the presence or absence of heat stress were compared (inserts; n ≥ 102 vacuoles). *P < 0.05. **(D)** Western blot analysis of Vba4-GFP, Cot1-GFP and Ypq1-GFP degradation before (0) and after (120 min) fusion of vacuoles isolated from wild type cells under standard conditions (CTL) or after heat stress (HS) in the presence or absence of fusion inhibitors rGdi1 and rGyp1-46 (F.I.). Pho8 is shown as a load control.

Since the ILF pathway relies on the vacuole fusion machinery for protein sorting and internalization (*McNally et al., 2017*), we reasoned that blocking the membrane fusion reaction would also inhibit protein sorting into the boundary membrane. To test this, we added the vacuole fusion inhibitors rGdi1 and rGyp1-46 (to block Rab-GTPase activation; *Brett et al., 2008*). As expected, blocking Rab activity prevented protein sorting (Figures 6A and B), internalization (Figure 6C), and proteolysis (Figure 6D) in response to heat stress, confirming that membrane fusion machinery, not ESCRTs, are required for observed polytopic protein degradation. In sum, we conclude that the ILF pathway, not vReD, is exclusively responsible for clearing misfolded vacuolar polytopic proteins, and thus is an important contributor to cellular protein quality control.

## DISCUSSION

### vReD and ILF pathways cooperate for lysosomal proteostasis

Despite being critical for lysosome physiology, it was unclear how lysosomal nutrient transporter proteins lifetimes were regulated until recently, when two processes were independently discovered: the ESCRT-dependent vReD pathway (*Li et al., 2015a*) and ESCRT-independent ILF pathway (*McNally et al., 2017*). However, if the two pathways function independently or cooperate to maintain or remodel the protein landscape on vacuolar lysosome membranes remained elusive. Herein, we find that they work together to control lysosomal proteostasis and begin to decipher their distinct contributions to this process: (1) Client proteins are shared by both pathways, as vReD client proteins Ypq1 and Cot1 are constitutively degraded by the ILF pathway (Figure 1). (2) Both pathways are responsible for constitutive turnover of vacuolar nutrient transporters, as we identified a new vReD client protein, Vba4, which is the first shown to be constitutively degraded by this pathway (Figures 2 and 3). (3) Similar to the ILF pathway, the vReD pathway responds to TOR activation suggesting that both may play a role in cellular metabolic homeostasis (Figure 4). (4) Only the ILF pathway appears to mediate quality control of lysosomal polytopic proteins, as all proteins studied are shunted into this pathway when misfolded upon heat stress (Figures 5 and 6). (5) The ILF pathway compensates for the loss of the vReD pathways when ESCRT function is impaired (Figures 2 and 3). Two questions immediately arise from these discoveries: What allows one pathway to compensate for the other? Why two pathways?

### What allows one pathway to compensate for the other?

Here we show that when components of the ESCRT machinery are deleted, the vReD pathway is blocked and the ILF pathway compensates for this loss by degrading Vba4 for example (Figure 2). We also find that the ILF pathway degrades internalized surface proteins that are normally packaged into intralumenal vesicles at the endosome by ESCRTs when MVB formation is impaired (*McNally and Brett, 2017*). These results provide important insight into the underlying mechanisms that may be shared. Fundamentally, both pathways require a mechanism to recognize and label client polytopic proteins, and selectively sort them into an area of membrane that is internalized into the organelle lumen for exposure to and degradation by acid hydrolases. These two pathways diverge at the sorting stage, where the sorting machinery is different (i.e. ESCRT-dependent versus ESCRT-independent), as are the membrane locations where sorting occurs and the mechanisms used for membrane severing required for internalization. Thus, we argue that they share the same labeling machinery. Selective client protein sorting by the vReD pathway requires protein ubiquitylation by E3-ubiquitin ligases and E4-adapter proteins (*Li et al., 2015a; Li et al., 2015b*). It is not surprising that the E3-ligase Rsp5 implicated in the vReD pathway also ubiquitylates surface proteins sorted into canonical MVB pathway, as both are ESCRT-dependent (*Horák, 2003; MacDonald et al., 2012*). This ligase is also important for mediating soluble protein labeling for degradation by the proteasome (*Glickman and Ciechanover, 2002*), highlighting that most degradation pathways share protein labeling machinery.

However, the basis of client protein labeling in the ILF pathway remains unknown. But given that it can degrade vReD client proteins in the absence of ESCRT function and all cellular degradation pathway seem to share protein labeling machinery, we speculate that the ILF pathway also recognizes and sorts client proteins ubiquitylated by E3-ligases and E4-adapter proteins. We have initiated studies to test this hypothesis and preliminary work indicates that the E4-adapter protein Ssh4 responsible for Ypq1 ubiquitylation for degradation by the vReD pathway (*Li et al., 2015a*), also mediates degradation of the copper-oxidase Fet5 by the ILF pathway (*E.K. McNally and C.L. Brett, unpublished results*). Thus, because they share the same protein labeling machinery, it seems that the ILF and vReD pathways coordinate functions with other cellular protein degradation pathways to restructure the cellular proteome, e.g. for metabolic reprogramming which requires remodeling the proteomes of lysosomes, mitochondria, peroxisomes, lipid droplets, ER, cytoplasm (*Lemus and Goder, 2014; Tasset and Cuervo, 2016*).

### Why two pathways?

There are two important distinctions between the vReD and ILF pathways that may explain why they co-exist: First, the vReD pathway permits vacuole polytopic protein recycling, whereby after sorting and delivery to the endosomes client proteins could avoid being packaged into ILVs by the ESCRT machinery. If so, they would remain on the MVB perimeter membrane, and upon MVB-vacuole fusion, be returned to the vacuole membrane. This is akin to surface receptor protein recycling, whereby after internalization, receptors on the endosome membrane can be packaged into ILVs for degradation or returned to the surface through retrograde trafficking (*Maxfield and McGraw, 2004*). Thus, we propose that the vReD pathway acts in a similar manner, to regulate levels of nutrient transporter proteins on vacuolar lysosome membranes. On the other hand, sorting into the ILF pathway ensures client protein degradation, as they are immediately exposed to lumenal proteases. This is based on the assumption that ILF-vacuole membrane “back” fusion does not occur, which is supported by the observations that GFP accumulates within the lumen in a diffuse pattern, not on puncta, over the course of the fusion reaction (Figures 1 – 6), and ILFs fragment and seem to dissolve quickly after being formed within the lumen (Videos 2 and 6; also see *McNally et al., 2017*). This explains why all misfolded proteins – which cannot be reused – are shunted into the ILF pathway for degradation (Figures 5 and 6).

The second distinction between the ILF and vReD pathways is organelle copy number: The vReD pathway can function when cells contain only a single vacuolar lysosome. However, because it requires merger of two organelle membranes, the ILF pathway functions only in cells with two or more vacuoles. Although this distinction seems trivial, it is worth noting that most micrographs presented in the reports by Emr and colleagues exclusively show cells containing single vacuoles (*Li et al., 2015a; Li et al., 2015b; Zhu et al., 2017*), which only permits analysis of the vReD pathway. Here we use an unbiased approach to study both pathways, by studying all cells in the population, i.e. those with single or multiple vacuoles. When conducting experiments, we did notice that GFP-positive puncta were more likely to be found in cells containing single vacuoles, suggesting that the vReD pathway may prevail in these cells. Whereas the ILF pathway would only contribute to vacuolar lysosome protein turnover in cells with multiple organelles. When considering this idea, it is important to note that metazoan cells contain hundreds to thousands of lysosomes (*Luzio et al., 2014*). These organelles are particularly mobile and frequently make contact often resulting in fusion. Each of these fusion events could accommodate protein turnover by the ILF pathway, which seems like a logical and efficient mechanism to maintain organelle homeostasis or accommodate membrane proteome remodeling.

Finally, it should be noted that Emr and colleagues who originally discovered the vReD pathway now propose that an endosomal intermediate may not be required (*Zhu et al., 2017*). Rather, the ESCRT machinery is thought to be recruited to microdomains on the vacuole membrane where it drives ILV formation directly into the lumen, similar to microautophagy (*Oku et al., 2017*). However, here we provide additional evidence to support their original model: First, we find that GFP-positive puncta continue to form, albeit less efficiently, when components of the ESCRT machinery are deleted (e.g. Figures 3A and 3B), suggesting that client proteins accumulate on endosomal intermediates adjacent to the vacuole membrane – presumably because the machinery that redistributes them on endosomes remains intact (i.e. retrograde vacuole-endosome trafficking; *Liu et al., 2012*) but here they cannot be packaged into ILVs by ESCRTs. Many of these GFP-positive puncta do not seem to contact the vacuole membrane, as originally reported (*Li et al., 2015a*), suggesting their presence on a separate endosome intermediate. Neither observation can explain the alternative, where we would expect ESCRT-dependent formation of GFP-labeled microdomains on the vacuole membrane to be observed as puncta exclusively located on the vacuole membrane and to be entirely abolished when ESCRT components are deleted.

Second, we find that protein degradation by the vReD pathway is blocked by protein inhibitors that target both endosome and vacuole membrane fusion in vitro (e.g. Vba4-GFP; Figure 6D). This finding is consistent with the original report (*Li et al., 2015a*) but contradicts the most recent report by Emr and colleagues that employs temperature-sensitive alleles of Vps18 and Vam7 to conclude that this fusion machinery is not required for protein degradation by the vReD pathway (*Zhu et al., 2017*). However, it is noteworthy that heat stress applied to suppress Vps18 or Vam7 also stimulates protein degradation by ESCRT–dependent and –independent processes (*Babst, 2014; McNally et al., 2017*; also see Figure 5 and 6). Furthermore, the approach used to stimulate client protein degradation, termed “RapiDeg”, requires use of rapamycin, an inhibitor of TOR signaling, which is an important mediator of cellular protein degradation (*Dobzinski et al., 2015; McNally et al., 2017;* also see Figures 4 and S2). Thus, these methods cause off-target effects that affect functions of the vReD and ILF pathways making it is difficult to interpret the results of these experiments. Adding further to this debate is the recent discovery that ESCRT-dependent microautophagy requires the vacuole fusion machinery for membrane scission (*Oku et al., 2017*). Small vesicles bud into the vacuole lumen from the tip of autophagic tubes (*Müller at al., 2000*) and resemble the products of the vReD pathway (*Zhu et al., 2017*). However, it is not clear whether a similar mechanism drives ILV formation by the vReD pathway, as Emr and colleagues do not show intermediates of vacuole ILV formation to support their hypothesis that the vesicles bud inwards from the vacuole membrane. Future studies will resolve this issue but regardless of outcome, it is clear that this process is distinct from the ESCRT-independent ILF pathway, whereby the products (ILFs) are obviously generated during homotypic vacuole fusion events within living cells (Videos 1-6; *McNally et al., 2017*).

### Physiological relevance

What roles do these pathways potentially play in lysosome and cell physiology? All proteins have finite lifetimes. Lysosomal transporter proteins are proposed to have shorter than average lifetimes because their lumenal faces are exposed to acid hydrolases making them particularly susceptible to damage, even in the presence of the glycocalyx (*Settembre et al., 2013*). In support of this idea, here we find that both the ILF and vReD pathways constitutively turnover nutrient transporters from the vacuolar lysosome membrane. Without such mechanisms in place, damaged proteins would accumulate, which is thought to permeabilize the lysosome membrane. This in turn facilitates release of lumenal hydrolases into the cytoplasm, which is catastrophic for the cell and implicated in cell death programs (*Aits and Jäättelä, 2013; Donida et al., 2017*). Here we find the ILF pathway clears and degrades misfolded lysosomal polytopic proteins and thus we speculate that its function is critical for preserving lysosome integrity necessary for cell survival or it may be inhibited during programmed cell death.

Beyond potential roles in organelle homeostasis, the protein landscape on the lysosome membrane defines organellar signaling properties (by possibly regulating levels of Ca^2+^ transporter or channels; *Perera and Zoncu, 2016*) and metabolic output (by changing the repertoire of nutrient transporters; *Lim and Zoncu, 2016*). This is akin to altering the expression profile of transporters or receptors on the plasma membrane in response to changes in nutrient levels or cellular signaling to mediate a myriad of physiological events. Here we show that the ILF and vReD pathways respond to similar stimuli to selectively downregulate some nutrient transporters and speculate that this alters organelle function for diverse cell physiology, including the cellular aging program that depends on changes in lysosome function (*Carmona-Gutierrez et al., 2016*). The machinery underlying both pathways and most client proteins are evolutionarily conserved, suggesting that they contribute to lysosome physiology in all eukaryotic cells. However, currently neither pathway has been shown to function in metazoan cells. But we hypothesize that they also determine lysosomal polytopic protein lifetimes and contribute to lysosome physiology in a similar manner, whereby they may also contribute to human lysosomal disorders linked to mutations in lysosomal transporters, e.g. the cholesterol transporter Ncr1 and Niemann-Pick type C disease (*Parenti et al., 2015*).

## EXPERIMENTAL PROCEDURES

### Yeast strains and reagents

All *Saccharomyces cerevisiae* strains used in this study are listed in Table S1. Biochemical reagents and yeast growth media were purchased from either Sigma-Aldrich, Invitrogen or BioShop Canada Inc. Proteins used include recombinant Gdi1 purified from bacterial cells using a calmodulin-binding peptide intein fusion system (*Brett and Merz, 2008*) and recombinant Gyp1-46 (the catalytic domain of the Rab-GTPase activating protein Gyp1) purified as previously described (*Eitzen et al., 2000*). Reagents used in fusion reactions were prepared in 20 mM Pipes-KOH, pH 6.8, and 200 mM sorbitol (Pipes Sorbitol buffer, PS).

### Fluorescence microscopy

Images were acquired with a Nikon Eclipse TiE inverted microscope equipped with a motorized laser TIRF illumination unit, Photometrics Evolve 512 EM-CCD camera, CFI ApoTIRF 1.49 NA 100× objective lens, and 488 nm (50 mW) solid-state laser operated with Nikon Elements software. Cross sectional images were recorded 1 μm into the sample and resulting micrographs were processed using ImageJ software (National Institutes of Health) and Adobe Photoshop CC. Images shown were adjusted for brightness and contrast, inverted and sharpened with an unsharp masking filter. Fluorescence intensity profiles of GFP fluorescence were generated using ImageJ software.

### Live cell microscopy

Live yeast cells were stained with FM4-64 to label vacuole membranes and prepared for imaging using a pulse-chase method as previously described (*Brett et al., 2008*). Where indicated, cells were incubated at 37 °C for 30 minutes (heat stress), with 100 μM cycloheximide at 30 °C for 90 minutes, or with 7 μM rapamycin at 30 °C for 90 minutes after FM4-64 staining. Time-lapse videos were acquired at 30°C using a Chamlide TC-N incubator (Live Cell Instruments) with cells plated on coverslips coated with concavalin-A (1 mg/ml in 50 mM HEPES, pH 7.5, 20 mM calcium acetate, 1 mM MnSO4). To assess how changing substrate levels affect polytopic protein sorting, yeast cells expressing Ypq1-GFP were incubated over night at 30 °C in YPD media supplemented with 125 μM lysine. Cells were back diluted in fresh YPD media supplemented with lysine and stained with FM4-64 for one hour at 30 °C. After two washes, cells were resuspended in SC media lacking lysine for 6 hours before collection and imaging. Yeast cells expressing Cot1-GFP were incubated over night at 30 °C in SC media. Cells were then back diluted in fresh SC media for FM4-64 staining (one hour at 30 °C), washed and resuspended in SC media supplemented with 2 mM ZnCl_2_ and grown for two hours at 30 °C prior to collection and imaging.

### Vacuole isolation and homotypic vacuole fusion

Vacuoles were isolated from yeast cells as previously described (*Haas, 1995*). Vacuolar membranes were labeled by treating vacuoles with 3 μM FM4-64 for 10 minutes at 27 °C. Fusion reactions were prepared by adding 6 μg of vacuoles isolated from GFP-expressing cells to standard fusion reaction buffer (containing 0.125 M KCl, 5 mM MgCl_2_, 1 mM ATP and 10 μM CoA in PS buffer) and then incubated at 27 °C for 60 minutes, unless otherwise noted, prior to being placed on ice for visualization by fluorescence microscopy. To block fusion in vitro, 3.2 μM rGyp1-46 or 4 μM rGdi1 (F.I.) were added to vacuole fusion reactions. Where indicated, vacuoles were pretreated with either heat stress (37 °C for 5 minutes), 100 μM cycloheximide (15 minutes at 27 °C), 7 μM rapamycin (15 minutes at 27 °C), 25 μM ZnCl_2_ (5 minutes at 27 °C) prior to being added to fusion reactions. To block effects caused by CHX, vacuoles were treated with 7 μM rapamycin prior to addition of cycloheximide (where indicated). Homotypic vacuole membrane fusion was assessed using a content-mixing assay based on a complementary, split β-lactamase approach (*Jun and Wickner, 2007; Karim et al., 2017*). For all concentrations of ZnCl2 tested, isolated vacuoles were pretreated with ZnCl2 for 5 minutes at 27 °C, then added to fusion reactions and further incubated for 90 minutes at 27 °C.

### Western blot analysis of isolated vacuoles

Fusion reactions were prepared as previously described using vacuoles isolated from yeast cells expressing a GFP-tagged vacuolar membrane protein. Where indicated, samples were pretreated with heat stress, cycloheximide, or rapamycin and with fusion inhibitors rGyp1-46 and rGdi1. Samples were incubated for 0 (ice), 30, 60, 90, or 120 minutes at 27 °C, placed on ice and then protease inhibitors (6.7 μM leupeptin, 33 μM pepstatin, 1 mM PMSF and 10.7 mM AEBSF), 1% DDM and 5X laemmli sample buffer were added. To solubilize membrane proteins but avoid aggregation, reactions were then incubated at 27 °C for 10 minutes prior to conducting SDS-PAGE. Nitrocellulose membranes were probed with anti-GFP (Abcam, Cambridge, UK) or anit-Pho8 (a kind gift from Alexey Merz, University of Washington, Seattle, USA) antibodies. All experiments shown were repeated at least three times. Blots were imaged by chemiluminescence using a GE Amersham Imager 600, and resulting images were prepared using Adobe Photoshop and Illustrator CC software.

### Western blot analysis of whole cell lysates

Yeast cell lysates were prepared as previously described (*Volland et al., 1994*). Briefly, extracts were prepared by collecting yeast cells at mid-log phase spinning 2,500 × g for 5 minutes. Cells were washed once with YPD, resuspended in fresh YPD and incubated at 30 °C for 90 minutes (control), at 37 °C for 90 minutes to induce heat stress, or at 30 °C for 120 minutes in the presence of 100 μM cycloheximide. After treatment, 5 OD_600_ units of cells were collected, pellets were resuspended in 0.5 ml lysis buffer (0.2 M NaOH, 0.2% β-mercaptoethanol) and incubated on ice for 10 minutes. Trichloroacetic acid was then added to a final concentration of 5% and followed by an additional 10 minutes on ice. Precipitates were then collected by centrifugation (12,000 × g for 5 minutes at 4 °C) and resuspended in 35 μl of dissociation buffer (4% SDS, 0.1 M Tris-HCl, pH 6.8, 4 mM EDTA, 20% glycerol, 2% β-mercaptoethanol and 0.02% bromophenol blue). 0.3 M Tris-HCl, pH 6.8 was added and samples were heated at 37 °C for 10 minutes. Samples were run on a 10% SDS-polyacrylamide gel and transferred to nitrocellulose membranes. Antibodies used for blotting were raised against GFP (B2; Santa Cruz Biotechnology, Inc.) or G6PDH (A9521; Sigma-Aldrich).

### Data analysis and presentation

To calculate the relative intensity of GFP fluorescence present in boundary membrane or vacuole lumen, we analyzed only docked vacuoles observed in fluorescence micrographs of vacuole fusion reactions (see *McNally et al., 2017*). GFP fluorescence intensities were measured using a ROI 4x4 pixels in diameter and acquiring a fluorescence value on the outer vacuole membrane, within the lumen and on the boundary membrane using ImageJ software. Fluorescence intensity values were correlated to individual vacuole size (boundary membrane length, vacuole surface area and circumference). Data are reported as mean ± SEM. Comparisons were calculated using Student two-tailed t-test, *P* values < 0.05 indicate significant differences (*). Quantitative data was acquired using ImageJ software and plotted using Prism GraphPad version 7.0 software. Figures were prepared using Adobe Illustrator CC software.

## AUTHOR CONTRIBUTIONS

C.L.B and E.K.M. conceived the project. E.K.M. performed experiments and prepared all data for publication. E.K.M. and C.L.B. wrote the paper.

## ACKNOWLEDGEMENTS

We thank Alexey J. Merz for providing antibodies. We also thank the Canadian Foundation for Innovation and Natural Sciences and Engineering Research Council of Canada for generous support of the Centre for Microscopy and Cellular Imaging at Concordia University. This work was supported by NSERC grants RGPIN/403537-2011 and RGPIN/2017-06652 to C.L.B.

---

**Table S1.**
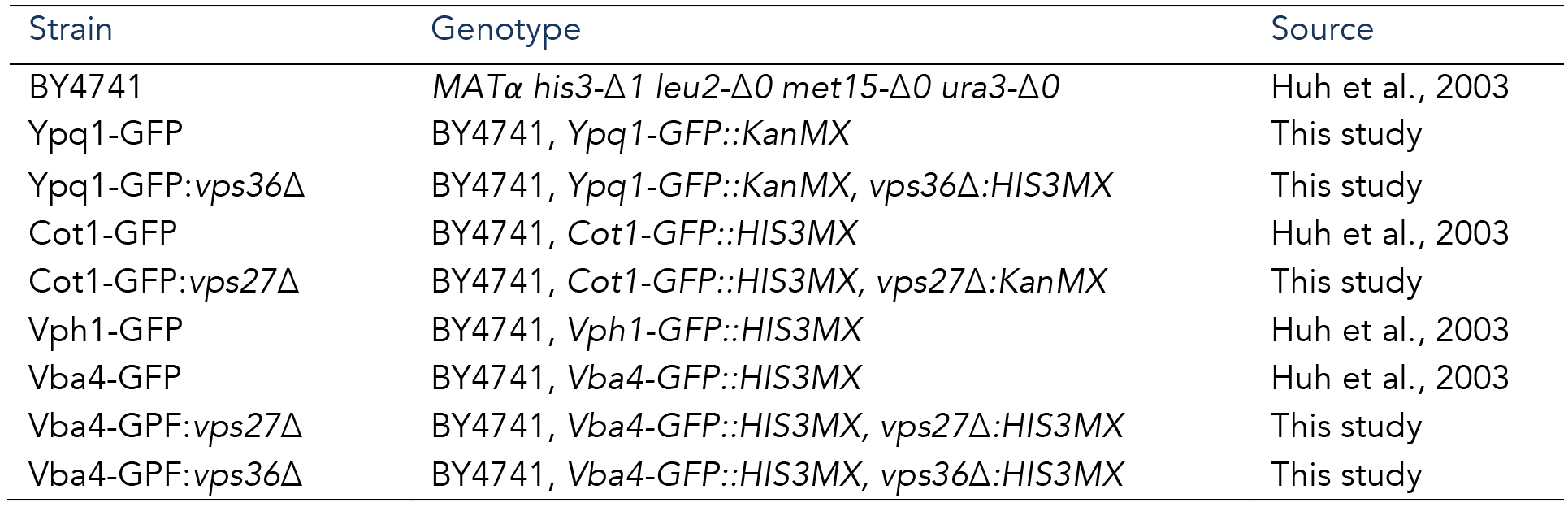
Yeast strains used in this study.

| Strain | Genotype | Source |
| --- | --- | --- |
| BY4741 | *MATα his3-Δ1 leu2-Δ0 met15-Δ0 ura3-Δ0* | Huh et al., 2003 |
| Ypq1-GFP | BY4741, *Ypq1-GFP::KanMX* | This study |
| Ypq1-GFP:*vps36Δ* | BY4741, *Ypq1-GFP::KanMX, vps36Δ:HIS3MX* | This study |
| Cot1-GFP | BY4741, *Cot1-GFP::HIS3MX* | Huh et al., 2003 |
| Cot1-GFP:*vps27Δ* | BY4741, *Cot1-GFP::HIS3MX, vps27Δ:KanMX* | This study |
| Vph1-GFP | BY4741, *Vph1-GFP::HIS3MX* | Huh et al., 2003 |
| Vba4-GFP | BY4741, *Vba4-GFP::HIS3MX* | Huh et al., 2003 |
| Vba4-GPF:*vps27Δ* | BY4741, *Vba4-GFP::HIS3MX, vps27Δ:HIS3MX* | This study |
| Vba4-GPF:*vps36Δ* | BY4741, *Vba4-GFP::HIS3MX, vps36Δ:HIS3MX* | This study |

**Figure S1, related to Figure 3.**
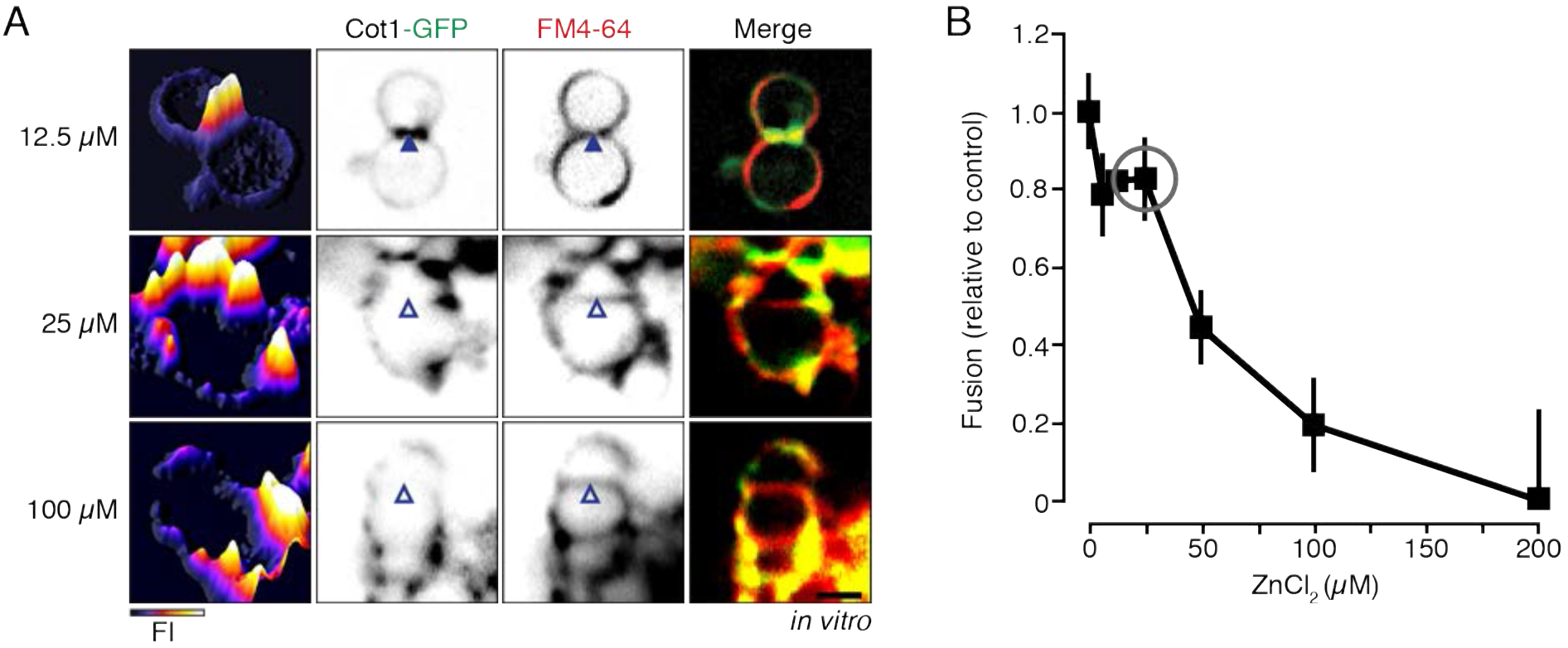
Conditions used to trigger Cot1-GFP degradation by the vReD pathway do not impair homotypic vacuole membrane fusion in vitro. **(A)** Fluorescence micrographs of isolated vacuoles from cells expressing Cot1-GFP pretreated with increasing concentrations of ZnCl_2_. **(B)** In vitro homotypic vacuole fusion of isolated vacuoles was measured after reactions were pretreated for 5 minutes at 27 °C with increasing concentrations of ZnCl_2_. Grey circle indicates ZnCl_2_ concentration (25 μM) used in all other experiments shown. Data was normalized to values obtained at 90 minutes under standard fusion conditions. Mean ± S.E.M. are shown (n = 2).

**Figure S2, related to Figure 4.**
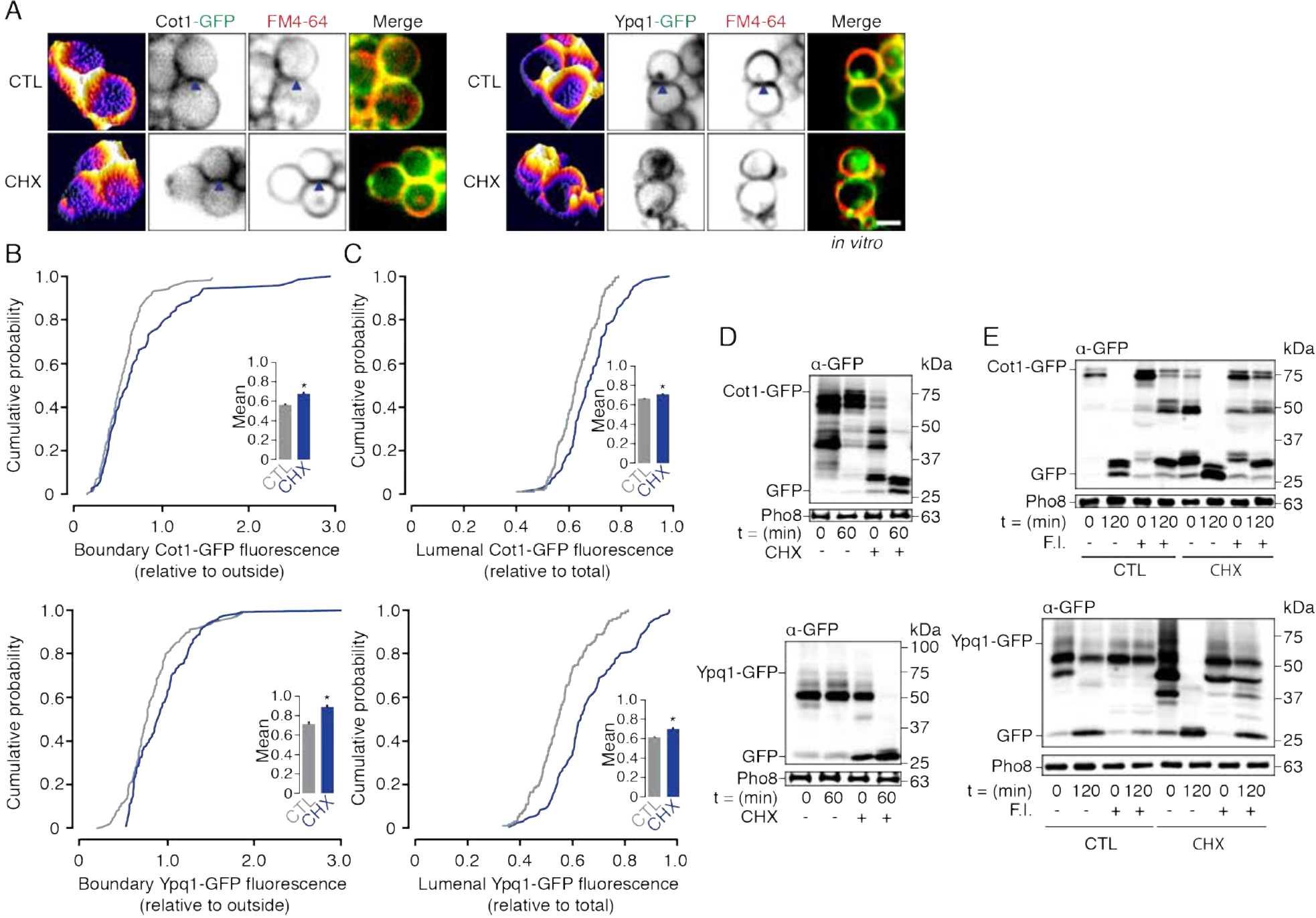
The ILF pathway degrades Cot1 and Ypq1 in response to TOR signaling in vitro. **(A)** Fluorescence micrographs of vacuoles isolated from wild type cells expressing Cot1-GFP or Ypq1-GFP stained with FM4-64 observed under standard fusion conditions (CTL) or after treatment with CHX. Boundary membranes containing GFP fluorescence are indicated by closed arrowheads. GFP fluorescence profiles are shown (left panels). Scale bar, 2 μm. Using micrographic data presented in **A**, we generated Cumulative probability curves of GFP fluorescence intensity within the boundary membrane **(B)** or vacuolar lumen **(C)**. Means (± S.E.M.) are plotted and values obtained in the presence or absence of CHX were compared (inserts; n ≥ 142). *P < 0.05. Western blot analysis of Cot1-GFP or Ypq1-GFP degradation before (0) and after (60 min) fusion of vacuoles isolated from wild type cells under standard conditions (CTL) or after treatment with cycloheximide (CHX; D), or in the absence or presence of fusion inhibitors rGdi1 and rGyp1-46 (F.I.; E). Pho8 is shown as a load control.

